# The recovery of parabolic avalanches in spatially subsampled neuronal networks at criticality

**DOI:** 10.1101/2024.02.26.582056

**Authors:** Keshav Srinivasan, Tiago L. Ribeiro, Patrick Kells, Dietmar Plenz

**Author notes:** Correspondence: Dietmar Plenz, Ph.D., Section on Critical Brain Dynamics, National Institute of Mental Health, Porter Neuroscience Research Center, Rm 3A-1000, 35 Convent Drive, Bethesda, MD 20892.

## Abstract

Scaling relationships are key in characterizing complex systems at criticality. In the brain, they are evident in neuronal avalanches—scale-invariant cascades of neuronal activity quantified by power laws. Avalanches manifest at the cellular level as cascades of neuronal groups that fire action potentials simultaneously. Such spatiotemporal synchronization is vital to theories on brain function yet avalanche synchronization is often underestimated when only a fraction of neurons is observed. Here, we investigate biases from fractional sampling within a balanced network of excitatory and inhibitory neurons with all-to-all connectivity and critical branching process dynamics. We focus on how mean avalanche size scales with avalanche duration. For parabolic avalanches, this scaling is quadratic, quantified by the scaling exponent, *χ* = 2, reflecting rapid spatial expansion of simultaneous neuronal firing over short durations. However, in networks sampled fractionally, *χ* is significantly lower. We demonstrate that applying temporal coarse-graining and increasing a minimum threshold for coincident firing restores *χ* = 2, even when as few as 0.1% of neurons are sampled. This correction crucially depends on the network being critical and fails for near sub- and supercritical branching dynamics. Using cellular 2-photon imaging, our approach robustly identifies *χ* = 2 over a wide parameter regime in ongoing neuronal activity from frontal cortex of awake mice. In contrast, the common ’crackling noise’ approach fails to determine *χ* under similar sampling conditions at criticality. Our findings overcome scaling bias from fractional sampling and demonstrate rapid, spatiotemporal synchronization of neuronal assemblies consistent with scale-invariant, parabolic avalanches at criticality.

## Introduction

Complex systems, composed of many local components or agents that interact weakly, often exhibit event cascades that cover a wide range of scales in both space and time. Such scale-invariant cascades, typically identified by power-law distributions in their duration and size – defined as the total activity observed during its lifetime, have been observed in many real systems, including solar flares^1^, earthquakes^2,3^, superconductors^4^, sandpiles^5^, forest fires^6^, and in the brain in the form of neuronal avalanches^7–9^. The scaling of such system cascades, specifically how their mean size grows with their duration, has been particularly informative as to the potential underlying dynamics of cascade generation. For example, if local system components activate independently at a constant rate, the mean size, <S>, of observed cascades will scale linearly with duration, T, exhibiting a scaling exponent of *χ* ≅ 1, i. e., temporally contiguous system events reflect a constant, yet random probability of their continuation. Values of *χ* ≅ 1 are often found near a 1^st^ order phase transition^10–12^.

Conversely, if cascades exhibit *χ* = 2, their mean growth is rapid and parabolic as can be found at the second-order phase transition of a high-dimensional critical branching process, supporting local system interactions^13^. In general, values of *χ* >>1 are of particular interest as they suggest the ability of a system to bind spatially distributed events rapidly and selectively into diverse internal system configurations which is generally considered beneficial for a complex adaptive system such as the brain (e. g.,^14,15^).

Here, we focus on neuronal avalanches, which were first shown to spontaneously emerge in isolated brain networks, i. e. acute cortex slices and long-term cortex cultures^7^. By measuring locally synchronized neuronal activity using the local field potential (LFP) and tracking the spatiotemporal spread of LFP cascades with microelectrode arrays, power-law distributions for both size, *S*, and duration, *T*, *P*(*S*) ∼ *S*^−*α*^ and *P*(*T*) ∼ *T*^−^*^β^* were revealed, with exponents α ≈ 3/2 and β ≈ 2 respectively, in addition to a critical branching parameter of σ ≅ 1, and significant correlation of neuronal activity within and between^16,17^ neuronal cascades (for review see^18^). These hallmarks of fast, neuronal synchronization in the form of avalanches were subsequently confirmed in the normal brain for ongoing and evoked LFP cascades in the cortex of nonhuman primates^19–22^ and in humans using magnetoencephalography^23,24^. Their sensitivity to even slight pharmacological alterations in the balance of excitation and inhibition^7,18,22,25^ and changes during pathological conditions such as epilepsy^26–29^ or changes in brain states such as sleep^30,31^ and sleep deprivation^32–35^, firmly indicate that these avalanches represent neuronal synchronization of cell assemblies in the normal brain.

To understand avalanche synchronization mechanistically, the contribution of single neuron firing to avalanches needs to be properly identified. This task is extremely challenging as studies in nonhuman primates and isolated slice preparations show that neurons participate sparsely and highly selectively even in spatially extended, large LFP avalanches^22^. Accurate reconstruction of avalanche synchronization from single neuron activity, therefore, requires the appropriate sampling of a significant portion of the brain network which poses an enormous technical challenge in neuroscience. Conversely, spatial subsampling, that is, sampling of only a fraction of neurons, is limited in identifying neuronal synchronization largely by failing to identify avalanche continuation in non-sampled neurons^36^. This underestimates avalanche duration and size, biasing outcomes towards expectations for decorrelated, random processes^9,37,38^. Accordingly, simulations of avalanche generating critical branching processes and fractional sampling of neurons reduces the expected growth in mean avalanche size with avalanche duration, *χ*, from the expected value of *χ* = 2, to a value of *χ* closer to 1 – 1.3^39–41^. Not surprisingly, estimates of *χ* ≅ 1.3 were reported for fractionally sampled neuronal spikes of potential avalanche activity *in vivo*^40,42–45^ and *in vitro*^46^.

If the exact start and end times of avalanches are known, for example in simulations under the assumption of ‘separation of time scales’, spatially subsampled avalanches can be easily corrected for by temporal integration thereby recovering the correct critical exponents in size and duration^41^ (for a review, see also^47^). Avalanche times, however, are largely unattainable in experimental data for which only a small fraction of neurons is observed and temporal integration eventually will degenerate avalanche scaling by indiscriminately concatenating successive avalanches^9^. Recently, Capek & Ribeiro et al.^9^ demonstrated that, even when the exact times of avalanches are not known, *χ* = 2 can be recovered for subsampled excitation-inhibition (E-I) balanced networks of integrate-and-fire neurons which are known to belong to the mean-field directed percolation (MF-DP) universality class^48,49^. This rescue requires (1) reducing uncorrelated neuronal action potential firing and (2) using temporal coarse-graining with an increase in minimal synchronization requirements to identify synchronized population activity in the network. Under these conditions, the authors could show *χ* = 2 *in vivo* 2-photon imaging recordings of cellular activity with high signal-to-noise ratio for sensory evoked and spontaneous spike avalanches in frontal and sensory cortices of awake mice in line with their simulations^9^. This is in line with a recent EEG study^50^ in humans that demonstrated *χ* = 2.

In the current work, we extend our previous investigations of rescuing *χ* in subsampled, critical networks. We systematically study the rescue of the scaling exponent *χ* to its critical value of 2 in a continuously driven, subsampled all-to-all DP-model^48^ for which we demonstrate the equivalence of fractional sampling and thresholding. Importantly, we show that the rescue requires the network to be very close to criticality and fails in the near sub- and supercritical regimes. We demonstrate the robust rescue of *χ* in the critical model over a domain covering 4 orders of magnitude in fractional sampling within the temporal coarse graining vs. thresholding parameter regime. Using this approach, we demonstrate the robust recovery of *χ* ≅ 2 in cellular 2-photon imaging data of the frontal cortex of awake mice over a large range of varying thresholds and recording conditions. We further show that, in contrast, the ‘crackling noise’ relationship and related measures which can predict *χ*^51–53^ fail under conditions of fractional sampling. Our findings demonstrate the pitfalls of assessing avalanche synchronization when only parts of a critical network are observed and demonstrate the regime of potential scaling rescue for synchronization in the form of parabolic avalanches under experimental conditions of fractional sampling.

## Results

Neuronal avalanches were studied using a critically balanced excitatory-inhibitory (E-I) model with all-to-all connectivity consisting of one million neurons driven externally using independent, stationary Poisson processes^48^ (**Fig. 1a**). The external driving triggered an average of 20 neurons to fire per time step, *Δt*, which intermittently induced spreading of neuronal activity in the network. Simulations spanned across 10^8^ time steps. An avalanche was defined as a series of contiguous time periods, *T*, of coincident firing within *Δt* surpassing a predetermined threshold, *θ*, over a specific number of time steps, *k* (**Fig. 1b, e**). Thus, our model-based approach was characterized by two basic parameters: *k·Δt* and *θ*, which were explored in the context of the subsampling fraction of observed neurons, *f*, ranging from 1 out of every 10,000 neurons observed (*f* = 0.01%) to all neurons observed (*f* = 100%). We identified the statistical distribution of sequence size, *S*, and duration, *T*, as a function of *θ*, *f*, and *k* and estimated the corresponding size exponent, *α*, and duration exponent, *β*, within the constraints of the lower and upper cut-offs (**Fig. 1c**; see Methods;^54,55^). In our critical model, these distributions take on the form of power laws at *k* = 1, f = 100% and *θ* = 100, the defining characteristics of avalanches^7^. The scaling exponent, *χ*, governs the growth in average avalanche size, 〈*S*〉, for a given duration, *T*,

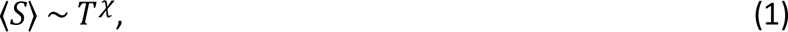

**Figure 1.**
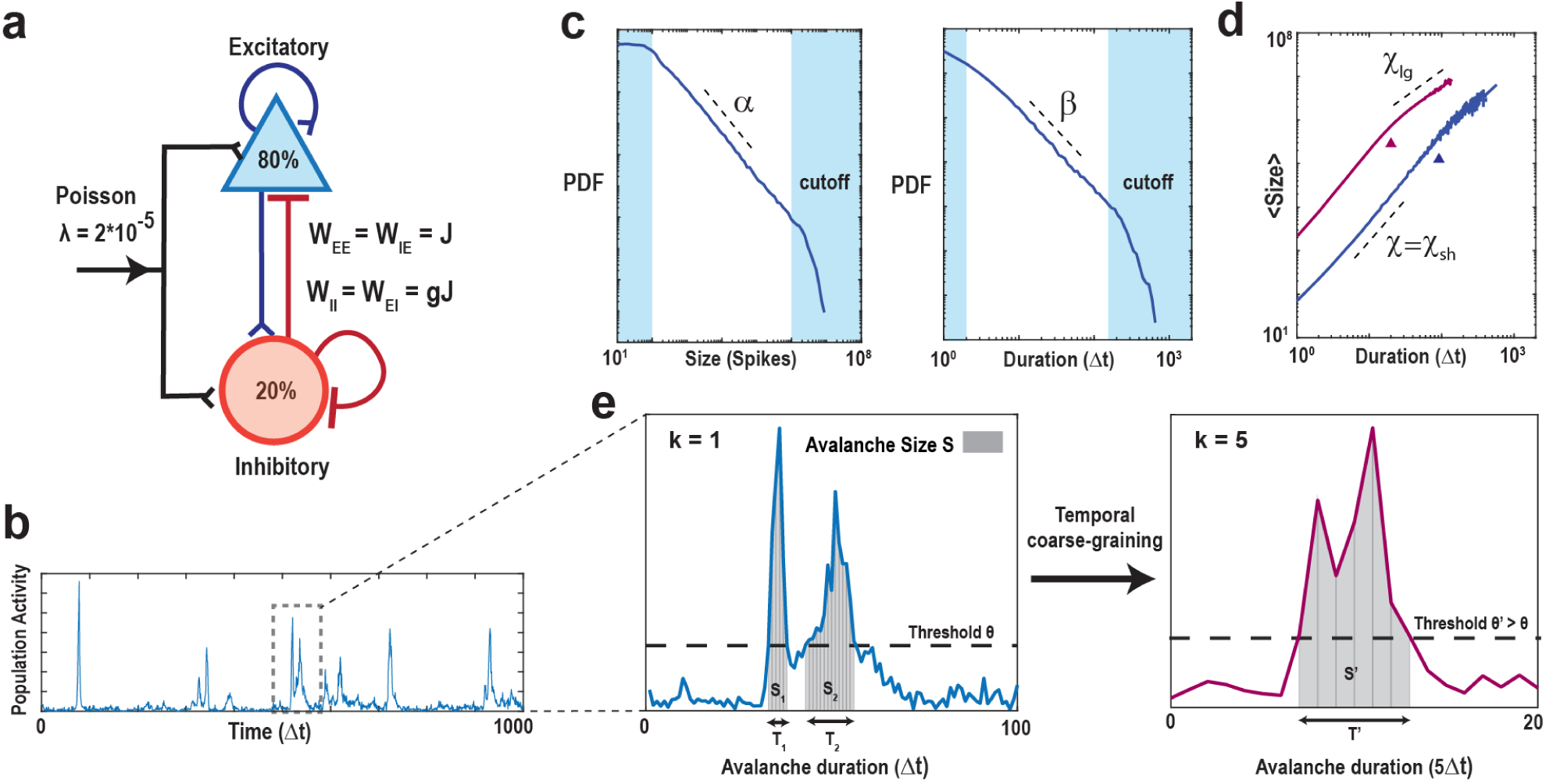
Schematics of the balanced E/I-model and identification of suprathreshold sequences in population spiking activity with respect to minimal coincident spiking threshold and temporal coarse graining. (**a**) Schematic of the neuronal network consisting of N = 10^6^ neurons (80% excitatory and 20% inhibitory) with a low external Poisson drive of rate *λ* = 2*10^-5^ per time step per neuron. The E/I balance is controlled by *g*, which scales the inhibitory weight matrices (W_II_, W_EI_) as a function of the excitatory weight matrices (W_EE_, W_IE_). If not otherwise stated, we set *g* = 3.5 to obtain critical dynamics (see Methods). (**b**) High variability in intermittent population activity characterizes critical dynamics. Snapshot of the summed neuronal spiking activity as a function of time. (**c**) Size and duration of avalanches in the critical model follow power-laws with corresponding exponent estimates *α* and *β*, respectively. Note that the external Poisson drive and the finite size of the network introduce lower and higher cut-offs, respectively (*shaded areas*). (**d**) Scaling of mean avalanche size as a function of duration also follows a power-law. *χ* is the estimate of the scaling exponent for short avalanches (below cut-off point, Δ). Avalanches within the upper cut-off (above cut-off point, Δ) exhibit a trivial scaling exponent close to 1, denoted as *χ_lg_*, that is largely independent of threshold and temporal coarse-graining and will not be considered further. *Purple*: Corresponding scaling relation and cut-off for temporal coarse-graining with k = 5 (see below). (**e**) Zoomed population activity from *B*. At the original temporal resolution *Δt* and given the coincident spiking threshold, *θ* (*left*), we can identify two sequences of suprathreshold activity, *S*_1_ and *S*_2_ with durations *T*_1_ and *T*_2_, respectively. Temporally coarse-graining the population activity (*right*; *k* = 5, binning the data into new bins of 5 time points) and increasing the threshold to *θ’ > θ*, absorbs *S*_1_ and *S*_2_ into a new suprathreshold activity period T’ with size *S*’ with corresponding change in scaling (see also d).

and accordingly, we estimated the scaling factor *χ* from the slope of the average avalanche size, 〈*S*〉, for a given avalanche duration, *T*, outside the minimal and maximal cut-off in logarithmic scales (**Fig. 1d**). This scaling factor was previously referred to as *χ_sh_* as it captures the scaling of short-duration avalanches that occur within the power law regime of the size and duration distribution (see also^9^). The fitting procedure (see Methods) also identifies a second slope, *χ_lg_*, for long-duration avalanches that occupy the cut-off regime of the corresponding size and duration distributions. As demonstrated previously and in line with expectations for uncorrelated activity^9^, *χ_lg_* remains close to 1 independent of thresholding and coarse-graining and this trivial scaling factor will not be considered further.

In critical systems, it has also been found that *χ* relates to the critical exponents *α*’ and *β*’ as

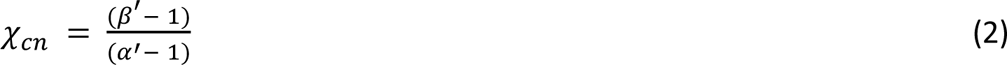

(’cn’ = crackling noise^51–53^). For comparison, we therefore calculated both *χ_cn_* and the ‘Deviation from Criticality Coefficient’ ^56^,

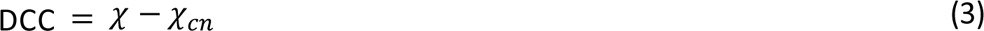

from our slope estimates *α* and *β*. This latter construct has increasingly been used as a distance measure from a critical point implying that the experimentally observed dynamics is close to criticality with low (< 0.2) DCC (e.g. ^44,57^), despite the fact that as one deviates from criticality the size and duration power laws break down, making the slope estimates *α* and *β*, and therefore the *χ_cn_* measure, unreliable.

### High coincident firing thresholds underestimate *χ* in the fully sampled model

In systems characterized by a clear separation of timescales and an absence of noise, the necessity for a coincident firing threshold diminishes. However, within real systems lacking such temporal separation and affected by significant intrinsic and measurement noise, the statistical analysis becomes intricately tied to our selection of threshold values. Setting the threshold too low poses the risk of incorporating substantial noise from the system, while setting it too high leads to data sparsity, resulting in the exclusion of a significant portion of the system’s activity. To emulate the noisy environment and absence of timescales separation between avalanche generation and propagation inherent in real systems, we introduce a minimal external Poisson drive in our simulations. This Poisson drive establishes a noise floor within the spiking activity, serving as the origin of avalanches within the system. This is clearly visible in the size distribution of avalanches, which exhibits a power law only for avalanche sizes of S_0_ > 50 – 100 spikes up to the finite-size of the network^48^ (*cf*. **Fig. 1c** *left*). To isolate avalanches above the noise floor (i. e., *S* > S_0_), we first studied the dependence of *α*, *β*, *χ* on *θ* and *k* in the fully sampled network (*f* = 100%; **Fig. 2**). In **Fig. 2a**, we show an overview in the form of color maps for all three parameters when increasing *θ*. At the highest temporal resolution (*k* = 1), we observed a rapid decrease in *χ* below its expected value of 2 as *θ* increases. This change largely follows the dependence of *β* while α remains close to 1.5, the latter being in line with findings from LFP avalanches in non-human primates^19^ (**Fig. 2b**). At high thresholds, *χ* tends to approach 1.3, a value commonly reported in experiments^39–45^, which is likely a consequence of subsampling biases inherent in these experimental settings. We conclude that high thresholds in weakly driven, noisy networks bias towards longer suprathreshold periods, significantly underestimating the scaling exponent in critical systems.

**Figure 2.**
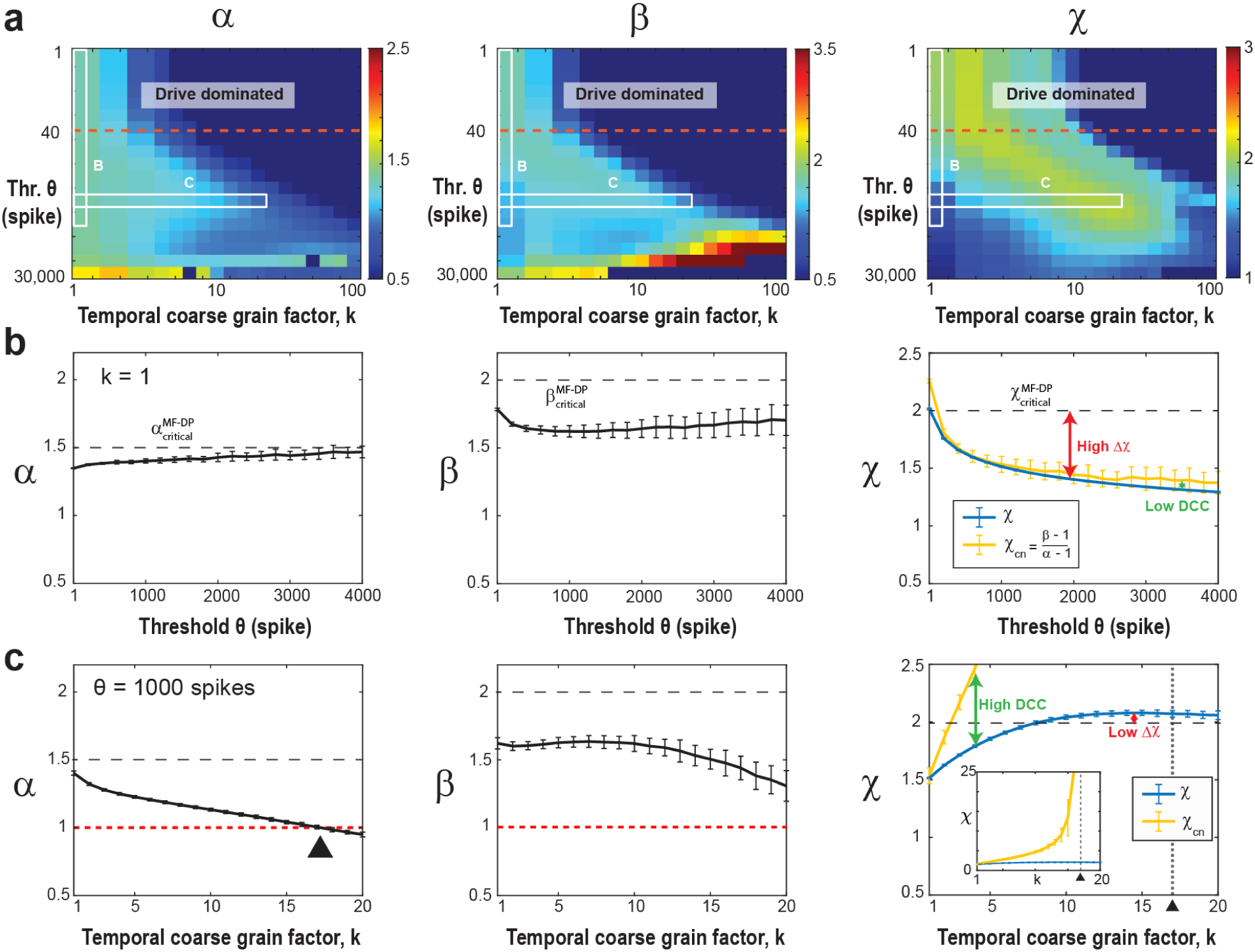
Increasing the threshold in the fully sampled model underestimates the scaling exponent, χ, which can be rescued by temporal coarse graining. (**a**) Consolidated view of the avalanche size (*α*), duration (*β*), and scaling (*χ*) exponents as a function of threshold *θ* and temporal coarse-graining factor *k* (*f* = 100%, fully sampled model). *Drive dominated*: Low threshold regime (*above the red broken line*) dominated by external Poisson drive. *White frames*: Parameter regions displayed in b and c. (**b**) At the highest temporal resolution (*k* = 1), high thresholds underestimate the scaling exponent, *χ*, as well as *χ_cn_*, with DCC remaining low. Size exponent, α (*left*), duration exponent, β (*middle*), scaling exponent, *χ*, and expected crackling noise relation, *χ_cn_*, (*right*) as a function of *θ*. (**c**) At highthreshold (*θ* =1000), temporal coarse graining recovers *χ*, but the *χ_cn_* exhibits a singularity leading to high DCC. Size exponent, α (*left*), duration exponent, β (*middle*), as a function of coarse graining factor, k, (*θ* = 1000). α passes through 1 (*red broken line*; *black triangle*), which causes a singularity in *χ_cn_*. (*right*). Note that the scaling exponent, *χ*, stabilizes to a value of 2 (*right, black dashed line*), whereas *χ_cn_* grows until it passes through a singularity at the temporal coarse graining value of *k* = 17 (*vertical broken line*; *triangle*; see *inset*).

A similar underestimation holds true for the *χ_cn_* value. Inserting the corresponding values of exponents extracted from our fully sampled system for small thresholds, we obtain *χ_cn_* ≅ 2 and DCC ≅ 0, in line with expectations for the mean-field directed percolation (MF-DP) universality class^58^.

When *θ* increases, *χ_cn_* drops significantly below 2, which is similar to our observation for *χ* and accordingly the DCC remains close to zero (see **Fig. 2a**). We attribute the small, yet consistently observed underestimation of *χ* with respect to *χ_cn_* (**Fig. 2b**) to the error-bounds of our fitting procedure. We conclude that avalanche scaling will be underestimated even in fully sampled networks if there are technical constraints on minimal coincident spike counts that require the use of high thresholds.

### Temporal coarse-graining rescues χ at high thresholds in the fully sampled model

Next, we show that temporal coarse graining rescues *χ* in the presence of a high threshold in fully sampled networks. This is shown in **Fig. 2c**, in which *χ* approaches 2 for *k* > 8 at *θ* = 1000 (see also corresponding projections in **Fig. 2a**). In contrast, the size and duration distributions exhibit a gradual reduction in steepness when *k* increases, due to concatenating subsequent cascades, in line with experimental findings^7,19,41,59^. In the model, this eventually causes *α* to cross the value of 1 (**Fig. 2c**), which forces *χ_cn_* to undergo a singularity (Equation 2; **Fig. 2c*, inset***). Accordingly, we find a rapid breakdown in the agreement between *χ* and *χ*_cn_ with increase in *k*, with diverging *DCC* (**Fig. 2c**). We conclude that while temporal coarse-graining can mitigate the bias introduced by thresholding in the fully sampled model, the *DCC* remains as an unreliable estimate for avalanche scaling even in fully sampled networks under high thresholding conditions.

### Rescue of χ = 2 is only seen in critical dynamics that is fractionally sampled and fails for slightly super- or sub-critical dynamics

We recently demonstrated that the rescue of the critical scaling exponent, *χ*, was exclusive to the critical model (*g_c_* = 3.5) and failed for subcritical conditions^9^. Here, we explored this dependency more systematically and show that small deviations from criticality by changing the E/I-balance parameter, *g* (see Methods), significantly impede rescue to the expected critical (MF-DP) value of 2. In **Fig. 3a**, we illustrate the maximum *χ* across various temporal coarse graining factors *k*, plotted against *g* for subsampled models (*f* = 0.1%) utilizing a 1-spike threshold (*θ* = 1). In subcritical models (*g* > 3.5), the restored scaling exponent swiftly deviates from its critical value of 2 towards 1. Conversely, in slightly supercritical models (3.45 < *g* < 3.5), *χ* also rapidly degrades to below 2. We note that for *g* < 3.45, persistent activity (shaded gray regions in **Fig. 3a**) obstructs the identification of avalanches at the specified threshold. This demonstration carried out for a critical branching model suggests that a successful rescue of the critical value of *χ* implies the dynamics to be at an E/I-balance of the order of 0.05 or 1.4% close to criticality. While further analytical studies are warranted to establish the rescue of *χ* as a key hallmark of critical dynamics, our findings suggest temporal coarse-graining combined with thresholding to recover the expected value *X* within a particular universality class might be a sensitive approach when confronted with fractionally sampled critical dynamics.

**Figure 3.**
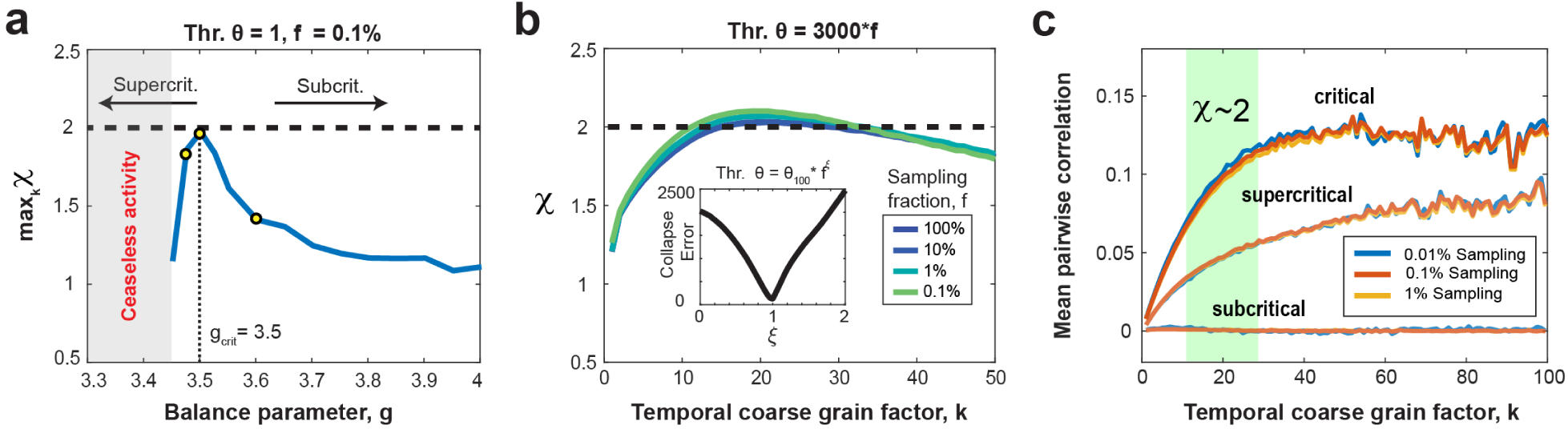
Temporal coarse graining at critical value of g: Rescue of scaling exponent, equivalence between threshold and fractional sampling, and increased mean pairwise neuron correlation. (**a**) The maximum scaling exponent across different temporal binning values as a function of the balance parameter, *g*, for a subsampled model (*f* = 0.1%) employing a 1-spike threshold. Notably, the retrieval of χ = 2 is exclusively observed in critical models (*g_c_*= 3.5), but promptly diminishes in slightly subcritical or supercritical models. In supercritical models, we obtain ceaseless activity (*shaded gray region*), precluding the identification of avalanches at the specified threshold. Yellow circles mark *g* values used in c. (**b**) Collapsed scaling exponent curves for different values of the sampling fraction, *f*, ranging from 100% to 0.1% for a threshold that is scaled by the sampling fraction as *θ = 3,000*f* (i.e., a threshold of 3,000 spikes for the fully sampled model translates to a threshold of 3 spikes for the 0.1% sampled model). *Inset*: The total collapse error as a function of collapse exponent, *ξ*. The error is taken over multiple different curves with *θ* ranging from 100 to 100,000. The error shows a clear minimum at *ξ* = 1, indicating the threshold can be scaled proportionally with the sampling fraction to obtain the best collapse. (**c**) The rescue of *χ* = 2 correlates with an increase of the mean delayed pair-wise correlation among neurons, which is independent of the fraction of neurons sampled. However, the temporal coarse graining regime of k for which *χ* = 2 (*green area*) is not predicted by this pair-wise correlation. For a supercritical (*g* = 3.475) network, the mean pairwise correlation is lower, but still increases with temporal coarse graining. For a subcritical network (*g* = 3.6), the mean delayed pair-wise correlation is close to 0.

### Scaling collapse demonstrates equivalence of thresholding and subsampling in the critically balanced all-to-all E/I-model

Both, the increase in coincident spiking threshold *θ* as well as a decrease in sampling fraction, *f,* underestimate the scaling exponent *χ* due to prematurely terminating avalanches by unobserved network activity. On the other hand, temporal coarse graining allows for concatenating sequential activity periods potentially recovering the proper duration for a subset of avalanches. We therefore examined the impact of temporal coarse graining on thresholding and fractional sampling. To achieve this, we first collapsed the *χ* vs. *k* curves over a wide range of *θ* (10^2^ –10^5^) by scaling *θ* with an exponent of *f*, *ξ* (see Methods). We found that the corresponding functions, obtained for different values of *f*, exhibited a minimum total collapse error for *ξ* = 1, demonstrating a linear relationship between *θ* and *f* in our all-to-all model. This implies that a fully sampled network (10^6^ neurons) with *θ* = 3,000 spikes is qualitatively equivalent to a 0.1% sampled model (1,000 neurons) with *θ* = 3 spikes (**Fig. 3b**). This equivalence between *θ* and *f* allows us to rescale the rescue of *χ* for different *f* in the corresponding (*k*, *θ*)-plane for ease of comparison (see **Fig. 4**).

**Figure 4.**
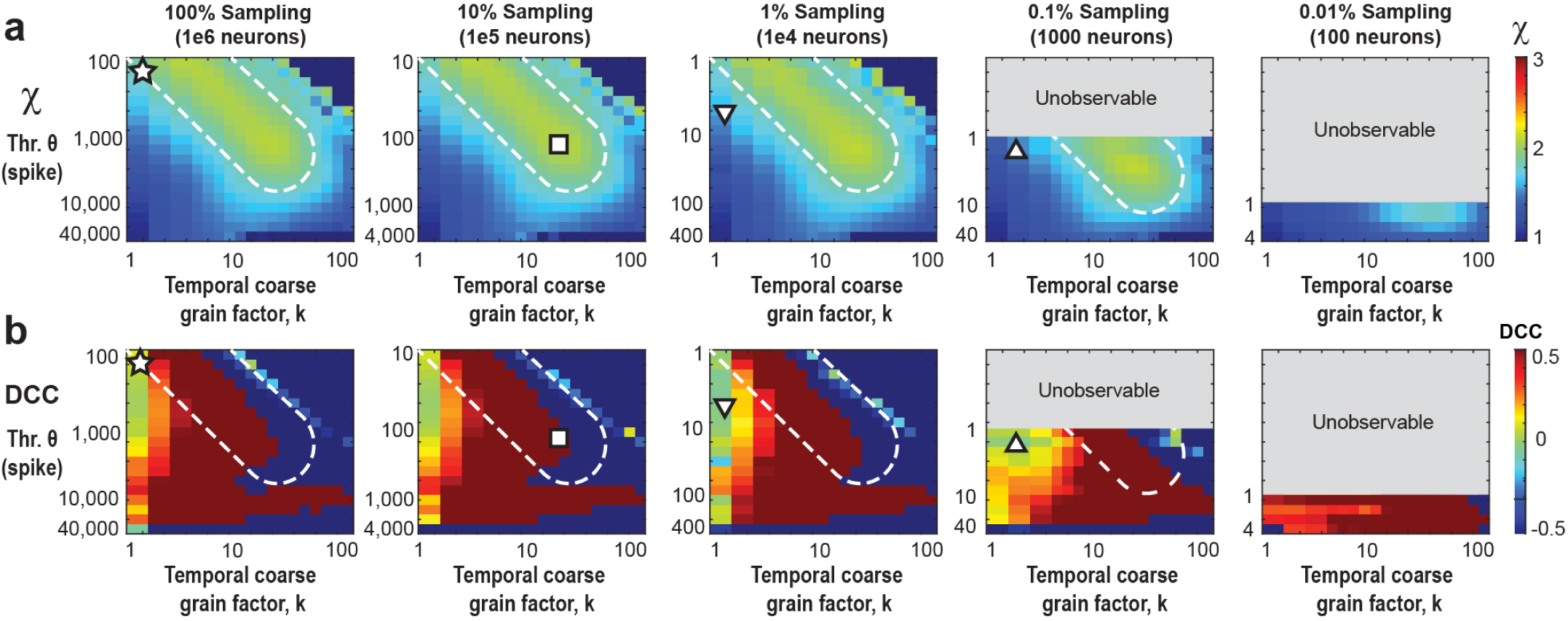
Temporal coarse-graining identifies a robust rescue regime for *χ* = 2 covering 0.1% - 100% fractional sampling at decreasing thresholding level, which is absent for the *DCC*. (**a**) Consolidated view of *χ* as a function of *θ* and *k* for different values of *f*. For *f* = 100%, *χ* ≅ 2 for low *θ* and *k* (*star*) but as we make the data sparser by increasing the threshold (or reducing the sampling), we need a higher coarse graining factor, *k*, to compensate and rescue *χ* back to 2 (*square*). *White dotted region*: visual guide for *χ* close to 2 for increasing *θ* and *k*. The grey parts of the plots for *f* = 0.1% and 0.01% respectively are unobservable parameter regions since they would require fractional thresholds, below the 1 spike minimum resolution of the model. At *f* = 0.1% sampling, *χ* can be rescued to a value of 2, but not for more severe subsampling, *f* = 0.01%. For triangles see b. (**b**) The *DCC* remains close to 0 only for low *k*, and quickly breaks down at higher *k* values. For the fully sampled model, there exists a region with the correct scaling exponent *χ* ≅ 2 as well as low *DCC* (*star* in *A* and *B*) However, as we move to lower sampling, this region of correspondence becomes more and more difficult to maintain. For f = 1% and 0.1%, regions at low k, and moderate threshold (or equivalently lower sampling) remain at low *DCC*, but underestimate the true scaling exponent, *χ* (*up* and *down triangles*, respectively).

### The rescue of *χ* = 2 indicates a recovery of mean pair-wise correlations

Subsampling as well as decorrelation of spiking activity have been shown to reduce the expected growth in mean avalanche size with avalanche duration from *χ* = 2, to a value of *χ* closer to 1 – 1.3^39–41^. We therefore studied how the mean delayed pair-wise correlation, 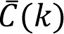, changes with temporal coarse graining, *k*, when recovering *χ* (see Methods). Temporal coarse graining increases 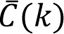 in the critical balanced E/I-network and saturates at high *k*, independent of the amount of fractional sampling (**Fig. 3c**). The increase in correlation did not predict the temporal coarse graining value of *k* at which *χ* = 2 and was not found in strongly subcritical networks, for which the average pairwise correlation remains near zero and independent of *k* (**Fig. 3c**; g = 3.75). In slightly supercritical models, the recovery of 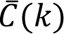 was significant but did not support *χ* = 2 for the critical model. Taken together, these findings demonstrate that *χ* = 2 identifies the expansion of coincidental firing among groups of neurons in the critical model and is not adequately captured by a particular value of 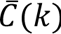, a measure limited to studying pairs of neurons.

### Subsampling and spike-resolution introduce unobservable regions of scaling exponents

By taking advantage of the trade-off between thresholding and sub-sampling (see Subsection above), we now can rescale our *k* vs. *θ* parameter space by *f· θ* to account for different fractional sampling. In the consolidated analysis (**Fig. 4**), we demonstrate reasonable agreement with the rescue of *χ* = 2 from 0.1% - 100% fractional sampling. When we increase the threshold (or equivalently, decrease *f*), we observe a deviation from *χ* = 2, yet the exponent can be reliably restored at higher values of *k* (**Fig. 4**, white broken lines). On the other hand, reducing *f* eventually introduces unobservable (*k*,*θ* )-regions which arise from the 1 spike resolution of the model excluding fractional thresholds. We note that even when sampling only 0.1% of the neurons, one can still recover *χ* = 2. However, this approach fails to simultaneously restore all exponents while maintaining the crackling-noise relationship (see **Fig. 2a** and Supplementary Fig. S1 online) because the point of concurrent rescue lies within the unobservable regions of parameter space. Decreasing sampling further to f = 0.01% (100 neurons out of 1 million neurons), prevents rescue of *χ* = 2.

Temporal coarse graining does not rescue the ‘crackling noise’ relationship of χ_cn_ (Equation 2). The DCC, introduced recently as the difference between the data-based value of χ (Equation 3) and *χ_cn_*should measure close to 0 if both estimates agree. As can be seen in **Figure 4**, co-regions with DCC = 0 and *χ* = 2 exist in the fully sampled system (**Fig. 4**; star). In contrast, no such co-alignment is found for subsampled systems and temporal coarse graining fails to establish a *DCC* = 0 regime for most values of *k* and *θ* (**Fig. 4**; square, triangles) and accordingly *DCC* = 0 does not reflect a proper rescue of the true *χ* = 2 under these conditions.

### Robust recovery of *χ* = 2 for parabolic avalanches in frontal cortex of awake mice

In the previous section, we demonstrated that temporal coarse-graining and thresholding effectively identify a robust subregion where the scaling exponent *χ* is rescued in the model. This rescue can be collapsed across different levels of fractional sampling. We now present similar findings from cellular 2-photon imaging of ongoing neuronal activity in the anterior cingulate cortex of awake mice (n = 17 experiments, each 30 minutes long; n = 5 mice; see Methods for details). The avalanche number distribution, obtained for each recording, follows a lognormal pattern as a function of the threshold (**Fig. 5a,d**; see also ^9^) and guides our analysis. In addition, this lognormal distribution varies with temporal coarse-graining and is thus a function of *k* (**Fig. 5a, d**). In a first approach, we pooled all data sets for a given absolute spike threshold (**Fig. 5a**), which revealed a consistent subregion of *χ* ≅ 2 for different spike density threshold levels (**Fig. 5b**). Specifically, we observe that the recovery of *χ* ≅ 2 shows a *θ* vs. *k* dependency, consistent with our simulations, whereas the DCC remains variable and noisy (**Fig. 5c**).

**Figure 5.**
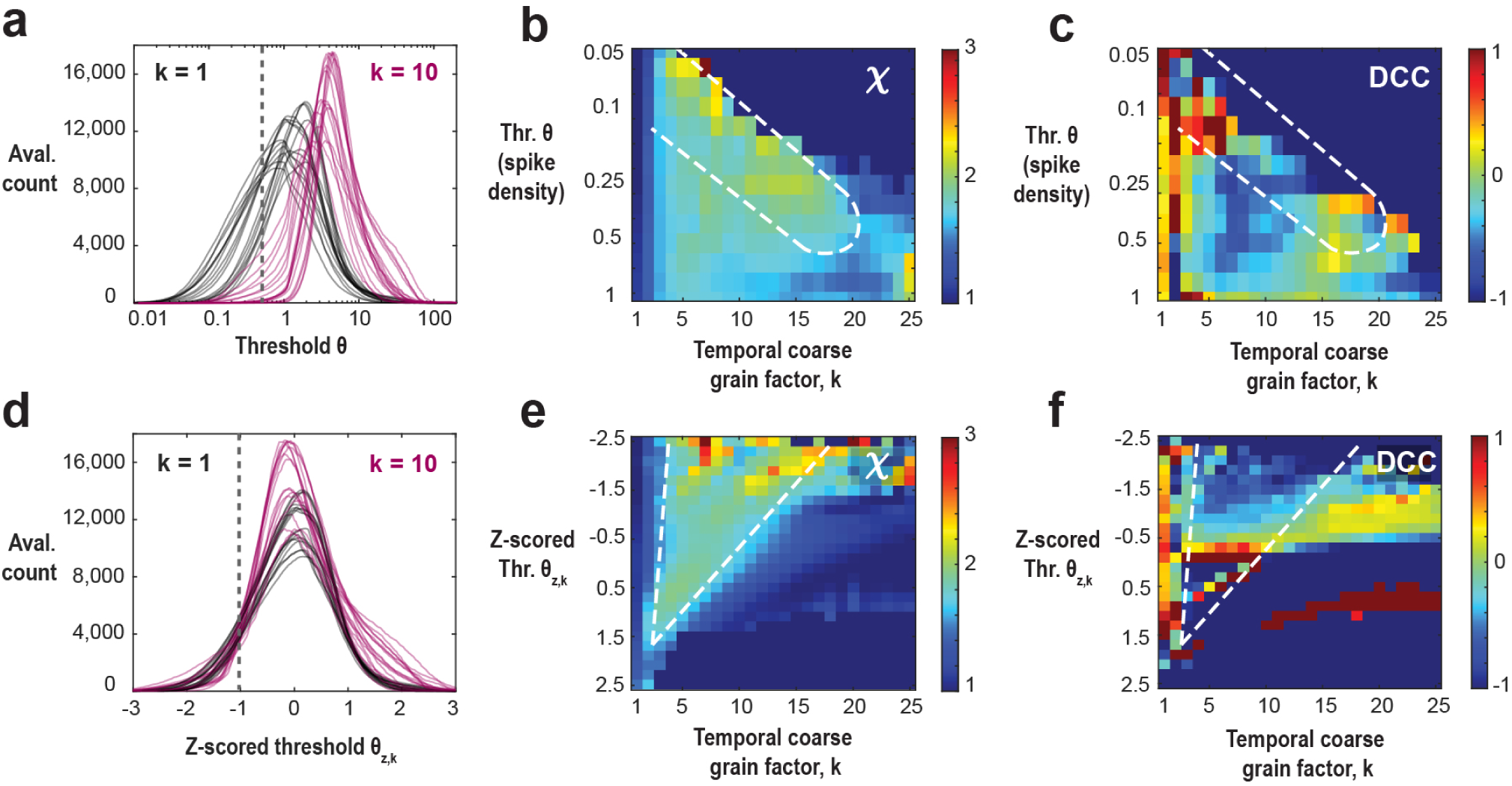
Parabolic avalanches in ongoing activity of frontal cortex exhibit threshold and temporal coarse-graining rescue of scaling exponent χ = 2 in line with critical model dynamics. **(a)** Lognormal distribution of avalanche number as a function of threshold for 17 different recordings (*n* = 5 mice; 2-photon imaging of ongoing activity in frontal cortex) at the original temporal resolution (*k*=1, *black*) and after temporal coarse graining (*k*=10, *purple*). Note the experimental variability at a given *k*, as well as the systematic shift in the distribution with change in *k*. A sample threshold (*θ, grey dotted*) shows how a given fixed threshold relates to these distributions. **(b)** Scaling exponent, *χ*, and **(c)** crackling noise deviation, DCC, as a function of temporal coarse grain factor, *k* and the spike density thresholds. The subregion in which *χ* ≅ 2 (*broken line*) is similar to the one identified in the critically balanced E-I model. At low thresholds, the scaling is rescued at smaller values of temporal coarse graining factor, k and at high thresholds, it is rescued at a larger value of *k*. Over this large range of parameters, the DCC is very noisy even a small change in the parameters can cause a large change in the DCC. **(d)** Z-scoring procedure for the lognormal distributions corrects for experimental variability as well as temporal coarse graining . A sample z-scored threshold (*θ_z,k_* , *grey dotted*) shows how a given z-scored threshold relates to these distributions. **(e)** Scaling exponent, *χ*, and (f) crackling noise deviation, DCC, as a function of temporal coarse grain factor, *k,* and the z-scored threshold at the given *k*, *θ_z,k_*. Figure D shows a robust rescue of *χ* ≅ 2 (*broken line*) for a large range of z-scored threshold values (-1.5 to 1.5). The range of k values over which *χ* ≅ 2 becomes smaller with an increase in the z-scored threshold. Like B, Figure E shows that the DCC remains unreliable and noisy even when the thresholds are z-scored and tuned for each *k*.

In a second approach, we z-scored the avalanche number distribution for each data set in order to account for experimental variability as well as for variable shifts induced by temporal coarse graining (**Fig. 5d**). **Figure 5e** shows that, regardless of the exact choice of z-scored threshold within a wide range (-2 to +1.5), *χ* is robustly recovered to approximately 2. The range of *k* values that recover *χ* decrases as the threshold increases. This robust recovery supports previous studies that adopted an avalanche-maximizing threshold (z-score = 0)^8^ or lower thresholds (z-score = -2)^9^. We note that at very low thresholds, noise becomes problematic, and at very high thresholds, data scarcity leads to poor statistics. In contrast, the DCC measure remains highly variable and unreliable even when the threshold is individually selected for each experiment and coarse-graining value (**Fig. 5f**).

## Discussion

The study of criticality in neuronal systems, particularly neuronal avalanches, has been extensively researched and debated (for review see^18,53,60–63^). This study addresses the intricacies and challenges of estimating scaling relationships in the brain, especially under conditions of fractional subsampling. A significant finding of our research is the ability to retrieve scaling exponents with fractional subsampling by employing temporal coarse graining and threshold adjustments. The sensitivity of this approach to deviations from criticality supports our method to study potential universality classes of criticality in real-world systems such as the brain. We demonstrate that temporal coarse-graining, combined with stricter criteria for coincident neuronal firing, allows for accurate estimation of the critical scaling exponent (*χ* = 2), even when observing as little as 0.1% of the network’s neurons. This suggests that it’s possible to accurately determine critical scaling characteristics despite heavy subsampling limitations. In contrast, previous studies on similar directed percolation (DP) models found a reduction in the scaling exponent to 1.3 with spatial subsampling of 10% and below^39^. Additionally, subsampling has been observed to falsely suggest population independence of correlations, impacting markers of criticality such as specific heat^64^. It’s important to note that our model, due to its all-to-all connectivity, does not allow for exploration of spatial correlation dependencies.

### Temporal coarse-graining when avalanche periods are unknown

Temporal coarse-graining has previously been suggested as a means to preserve the size exponents of avalanches in subsampled systems, assuming an infinite separation of time scales^41^. This assumption would theoretically allow researchers to distinguish sequentially occurring avalanches, a scenario which may not hold in natural systems. In contrast, our analysis shows that temporal coarse-graining can effectively maintain the scaling exponent in models lacking a clear separation of time scales, even with fractional sampling as low as 0.1%. This suggests its utility in analyzing highly subsampled experimental data within the directed percolation (DP) class. Our approach introduces a threshold above the Poisson noise floor to approximate the separation of successive avalanches, revealing a trade-off between subsampling intensity and the threshold level. In our all-to-all model, increasing the threshold of the fully sampled network mimics the effects of subsampling combined with a lower threshold, illustrating the interaction between these factors and their collective impact on criticality evaluations. Leveraging this trade-off, we apply our methodology to cellular-resolution 2-photon imaging data from the anterior cingulate cortex of awake mice, consistently recovering the scaling exponent to a value of 2, in line with expectations for a critical branching process. Additionally, new avalanche detection methods that utilize continuous time models have been developed, though their effectiveness compared to traditional methods under experimental and subsampling conditions remains to be fully evaluated^65^.

Our recovery of the scaling exponent *χ* ≈ 2 in ongoing neuronal spike synchronization of the awake mouse frontal cortex provides compelling evidence for the critical role of *χ* ≈ 2 in capturing essential features of neuronal avalanches (see also^9,20^). For instance, models with slow latent variables can generate heavy-tailed or even power-law distributions of neuronal activity size and duration, but they typically show *χ* ≈ 1, suggesting a lack of correlated activity^66–68^. While the critical branching process used here achieves *χ* ≈ 2, it does not account for temporal correlations among successive avalanches (e. g.^16–18,69^), limiting its ability to fully model neuronal avalanches in the brain.

It’s important to distinguish temporal coarse-graining from renormalization procedures. While renormalization in neuronal avalanches can preserve estimates of maximal mutual information during spatial coarse-graining under complex LFP sampling conditions^70^, our focus here is on addressing stochastic subsampling bias through temporal coarse-graining, as detailed previously^9^.

### Limits in rescuing the scaling exponent under subsampling conditions

Our simulations also clearly show that subsampling in discrete models renders certain parts of the phase space inaccessible (see **Fig. 4**), preventing the recovery of the scaling exponent under heavy subsampling (*f* << 0.1%). A similar limitation exist for temporal subsampling, especially when the required precision exceeds the system’s resolution^39^. Properly addressing these spatial and temporal constraints is crucial for the analysis of experimental data with limitations in both resolution domains. Further limitations arise with extensive subsampling of network topologies, particularly when estimating heavy-tailed or scale-free distributions^71–73^.

While under-sampled critical branching processes have been explored in extracellular spike recordings with microelectrode arrays in the anesthetized rat^36^, these limitations shouldn’t overshadow our findings’ potential. Careful consideration of experimental parameters can ensure feasible and informative criticality assessments, providing valuable insights into the core mechanisms of neural synchronization.

### The DCC and crackling noise relationship fails to recover the scaling exponent under subsampling conditions

The Deviation from Criticality Coefficient (DCC) approach^40,44,56^ has gained popularity for quantifying criticality based on the “crackling noise relationship”^51–53^. However, concerns have been raised about its ability to identify critical dynamics^37,68^ given that random processes can exhibit a DCC close to 0 thus limiting general claims of this approach to identify critical dynamics even in fully sampled systems. For instance, an unbiased random walk will produce excursions above the mean that are power-law distributed in sizes, with exponent 1.5, durations, with exponent 1.33 and scaling with exponent 1.5, which result in the random walk obeying the crackling noise relationship (*DCC* = 0), despite not being a critical system. Our study demonstrates several limitations of the DCC approach, particularly under subsampling conditions: (1) DCC estimates become unreliable with limited data, (2) the DCC is highly sensitive to temporal coarsening, leading to noisy results, (3) the DCC metric has a singularity under fractional sampling, making it a questionable marker for criticality. Furthermore, the DCC approach assumes power-law slopes obtained from size and duration distributions directly translate to critical exponents. This is flawed, as analysis parameters like temporal resolution and spatial sampling distance systematically influence these slopes^7,19^. These factors must be considered when interpreting criticality from experimental data using the DCC approach.

### The sensitivity of temporal coarse-graining approach to thresholds in avalanche size definition

Our analysis emphasizes the scaling exponent’s sensitivity to synchronization thresholds. As the threshold increases, we observe a noticeable deviation of the scaling exponent and the crackling noise ratio (*χ_cn_*) from the expected value of 2 (see also^13^). This sensitivity highlights the importance of careful consideration when choosing thresholding strategies in experimental data. Thresholding of time-series and ignoring the subthreshold contribution during suprathreshold periods can lead to a change in the avalanche size exponent itself as demonstrated in simulations of critical dynamics^13^. Capek, Ribeiro et al.^9^ demonstrated experimentally that including or excluding the subthreshold regime for size estimates provides narrow upper and lower bounds for *χ* that center around 2. Ideally, threshold selection should strike a balance between preserving genuine synchronization events and minimizing noise, especially when accurate scaling exponent estimates are crucial.

### Synchronization, coincident firing and delayed neuronal correlations

While traditional studies of spatiotemporal synchronization in neural systems have focused on phase-locked oscillations, our system operates in a unique scenario—one devoid of a dominant frequency. This lack of a dominant frequency becomes particularly important when considering ’noise correlations,’ which encapsulate the intrinsic non-stimulus-induced correlation structure among neurons. These correlations exert a notable influence on the decoding capabilities of neuronal populations, constituting a pivotal aspect of neural information processing^74–76^.

In this study, we investigate scale-free neuronal avalanches as a context for exploring neuronal synchrony and criticality. Avalanches offer a powerful tool for understanding neuronal synchronization, from local clusters to the entire brain. As we’ve shown, analyzing avalanches depends crucially on choosing thresholds and temporal resolution. This dependence is similar to the analysis of spatial and temporal neuronal correlations, which are heavily influenced by their respective coarse-graining^77,78^. Interestingly, as shown in Figure 3, restoring the scaling exponent *χ* to its critical value of 2 doesn’t directly correspond to the recovery in pairwise correlations. This emphasizes that *χ* at 2 signifies a critical system that describes the coordination among many neurons, not just an average estimate of correlation among pairs of neurons.

Our findings have implications for both neuroscience and complex systems research. Neuronal avalanches are valuable markers of synchronization in the brain. Precise scaling exponent estimation, even with limited data, is crucial for understanding mechanisms of neural synchronization in healthy and diseased brains. This supports the idea that the brain operates within a critical regime, aligning with the concept of self-organized criticality observed in many complex systems.

## Methods

### Model Topology

In this work, we employed the model originally proposed by Girardi-Schappo et al.^48^ and subsequently modified in Capek & Ribeiro et al.^9^. This model represents a critically balanced system of stochastic integrate-and-fire (IF) neurons, where the excitatory and inhibitory neurons maintain an equilibrium. Our neuronal network consisted of *N* = 10^6^ stochastic, non-leaky IF neurons, with a fully connected architecture. To reflect the prevalence of excitatory neurons in the cortex, we set the ratio of excitatory to inhibitory neurons to 4:1. The connectivity matrix, denoted as W, was initialized in a manner such that all outgoing synapses from excitatory neurons had equal strength, with W_EE_ = W_IE_ = *J*. Similarly, the inhibitory neurons had equal strength for their outgoing synapses, with W_II_ = W_EI_ = -*gJ*. Here, the balance parameter, *g*, plays a crucial role in tuning the model towards an E/I-balanced state exhibiting critical dynamics. For our network, the critical value of *g*, *g_c_*, was determined to be 3.5. When *g* exceeds *g_c_*, the network shifts towards an inhibition-dominated regime with subcritical dynamics, whereas values of *g* below *g_c_* cause the network to exhibit an excitation-dominated regime with supercritical dynamics.

### Model Dynamics

Each neuron in the model was described by two variables. The Boolean variable, X, indicated whether a neuron was firing or in a quiescent state at time t. Specifically, X(t) = 1 represented firing, while X(t) = 0 denoted quiescence at time t. The membrane potential, V, influenced the firing probability and evolved according to the following equation:

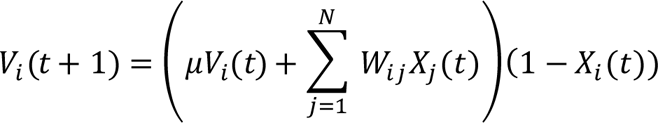

In the equation, μ represented the leakage parameter, which we set to 0 (non-leaky) for our simulations. The term (1 – X_i_(t)) introduced an absolute refractory period of *Δt*, allowing the voltage to reset after a spike occurred (when X_i_(t) = 1). Additionally, the probability of a neuron firing linearly increased with V and was given by:

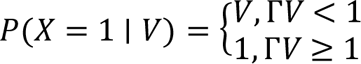

Here, Γ represented the neuronal gain, which we fixed at 1 for simplicity and supports probabilistic behavior. Notably, this network exhibited significant non-conservativeness, with energy dissipation occurring through inhibition and spike collision or spike coalescence^79,80^.

Additionally, we note that this model includes an absorbing quiescent state, and therefore, the network activity is initiated by very low-rate independent Poisson processes applied to each neuron. Specifically, the Poisson process triggers approximately ∼20 neurons (0.002%) on average per time step, which is negligible compared to the system size of 10^6^ neurons.

### Avalanche Statistics

To comprehensively investigate the resulting population activity, we run 100 simulations of the network over an extended duration of 10^6^ time steps for a total of 10^8^ time steps. For the case of full sampling, we compute the sum of activity across the entire population of *N* = 10^6^ neurons. However, to explore various sampling scenarios, we consider a random subset of *N*\**f* neurons, where *f* represents the sampling fraction. Following this, a thresholding procedure is applied to the time-series data, setting all neuronal activity below the threshold value *θ* to 0. We have the option to analyze the data within the thresholded time-series in their original form or subject them to temporal coarse graining. When employing coarse graining, each generation represents the sum of activity over *k* consecutive time bins, with *k* representing the coarse graining factor. This approach enables us to investigate network dynamics across different temporal scales and analyze resulting patterns. Consequently, we collect the sizes and durations of contiguous epochs from the time-series and examine their distributions.

### Power-law fitting

In the context of an ideal, infinitely sized critical system, the distribution of size, duration, and average size as a function of duration adheres to perfect power laws. However, it is important to consider statistical anomalies, finite size effects, and the influence of the low Poisson drive in our simulations when analyzing these distributions. Therefore, a cautious approach is required for their analysis within the model. In the average size versus duration analyses, we observed that the distributions could be reasonably approximated by a double power law. For this purpose, we employ the fitting function introduced by Capek & Ribeiro et al.^9^:

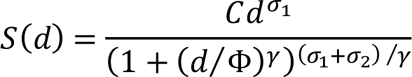

This function exhibits an initial slope σ_1_ that transitions to a second slope σ_2_ around the point *d* = *Φ*. The parameter γ governs the abruptness of this transition and has been set to 4 for all the analysis presented. The remaining parameters were allowed to adjust to the data, and all fits were performed in log-space. In all plots presented in this paper, σ_1_ is employed as the representative slope, while σ_2_, which is typically associated with finite-size effects, is not discussed in this context.

We use the cutoff, *Φ*, that we obtain from the scaling curves to analyze the individual size and duration distributions. The size exponent, *α*, is the slope of the size distribution in log-scale with a lower cutoff which is safely set to be greater than the Poisson noise level (>100) and an upper cutoff equal to the average size corresponding to the elbow point, *Φ*. For the duration exponent, *β*, we do a similar analysis, with a lower duration cutoff of 3 time-bins and an upper cutoff equal to *Φ*. However, for the sake of simplicity and due to the small variability of the fitting procedure for the experimental data, we choose to utilize only the first 4 generations to calculate estimates for all the exponents for Figure 5. Furthermore, we only consider exponents for parameters which have at least one avalanche that exceeds 20 generations and cases that do not satisfy this criterion are ignored.

### Collapse exponent of threshold and fractional sampling

In our analysis of the effects of temporal coarse graining on thresholding and fractional sampling, we collapsed the *χ* vs. *k* curves for *k* within the range 1 – 100. In the fully (100%) sampled model, we examined various thresholds (θ_100_) ranging from 10^2^ to 10^5^. In the case of fractionally sampled models, the threshold was adjusted proportionally to an exponent, *ξ*, of the sampling fraction, *f*, according to the equation θ = θ_100_ ∗ *f*^ξ^. For each ξ value spanning from 0 to 2 in steps of 0.1, we computed the collapse error of the five curves representing sampling rates from 0.01% to 100% of *χ* vs. *k*. The collapse exponent *ξ* is determined by the minimum collapse error and was found to be close to 1.

### Correlation analysis

In our examination of the mean delayed pairwise correlation within the system, we focus on a smaller temporal segment of the simulated network (consisting of 10,000 time steps) to facilitate analysis of large neuron raster datasets. For each neuron pair (designated as an ordered pair [i,j]) within the network, we compute the time-lagged correlation between the activity of neuron *i* and that of neuron *j*, shifted by one time step, *kΔt*, *C*_*ij*_ (kΔt). The mean delayed correlation, 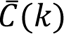, of the system is then determined as the average correlation coefficient computed for all such neuron pairs. Hence 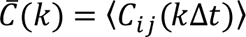. This correlation analysis spans temporal bins, *k*, ranging from 1 to 100 and spatial sampling values from 0.01% up to 1%. Due to the quadratic increase in duration and memory requirements with system size, higher sampling (10% sampling or more) values become impractical for storing raster data and conducting pairwise analyses.

### Ongoing activity in contralateral ACC/mPFC monitored with jRGECO1a

All experiments have been previously reported in Capek & Ribeiro et al.^9^ and are available online (see Data Availability). No additional experiments were conducted for the present study. In short, mice (C57BL/6; Jackson Laboratories; age >6 weeks) were injected with a viral construct to express jRGECO1a in cortical neurons using Syn promotor. Chronic 2PI started after ∼2 weeks in the contralateral ACC/mPFC at an estimated depth of ∼150– 300 μm using a microprism. Recordings were collected over the course of several days from n = 5 mice (3 males, 2 females; age 8–20 weeks) with jRGECO1a expression (n = 17 recordings; 30 min each). Recordings were conducted over the course of several weeks and analyzed separately for each mouse. 2PI images were acquired by a scanning microscope (Bergamo II series, B248, Thorlabs Inc.) coupled to a pulsed femtosecond Ti:Sapphire 2-photon laser (Chameleon Discovery NX, Coherent Inc.). The wavelength was tuned to 1120 nm to excite jRGECO1a. The field of view was ∼450 um x 450 μm. Imaging frames of 512 x 512 pixels were acquired at 45.527 Hz by bidirectional scanning of a 12 kHz Galvo-resonant scanner. The average power for imaging was <70 mW, measured at the sample. Obtained tif-movies in uint16 format were rigid motion corrected via the python-based software package suite2p^81^. Registered images were denoised using machine-learning-based, DeepInterpolation^82^ and then semi-automatically processed by suite2p for ROI selection and fluorescence signal extraction. For each labeled neuron, raw soma and neuropil fluorescence signals over time were extracted for each ROI. Spiking probabilities were obtained from neuropil corrected fluorescence traces (F_corrected = F_ROI – 0.7*F_neuropil) via MLspike (https://github.com/MLspike) by utilizing its autocalibration feature to obtain unitary spike event amplitude, decay time, and channel noise for individual ROIs.

## Acknowledgements

This research was supported by the Division of the Intramural Research Program (DIRP) of the National Institute of Mental Health (NIMH), USA, ZIAMH002797, ZIAMH002971, and the BRAIN initiative Grant U19 NS107464-01 to D.P. This research utilized the supercomputing resources of the National Institutes of Health (NIH, USA; Biowulf, http://hpc.nih.gov ) and the University of Maryland (UMD College Park, USA, https://hpcc.umd.edu/hpcc/dt2.html ).

## Author Contributions

K.S., T.L.R., D. P. conceived the study, K.S. did the simulations, P.K. provided the experimental data and processing, K.S., T.L.R., D. P. wrote the manuscript.

## Data Availability

The preprocessed imaging data used in this study are available in the general repository Zenodo using the following access https://doi.org/10.5281/zenodo.7703224 under ‘jRGECO1a-ongoing’.

## Code Availability

The code used to generate the model data is available at https://github.com/PlenzLab/ParabolicAvalanches/.

## Supplementary Information

**Suppl. Fig. S1.**
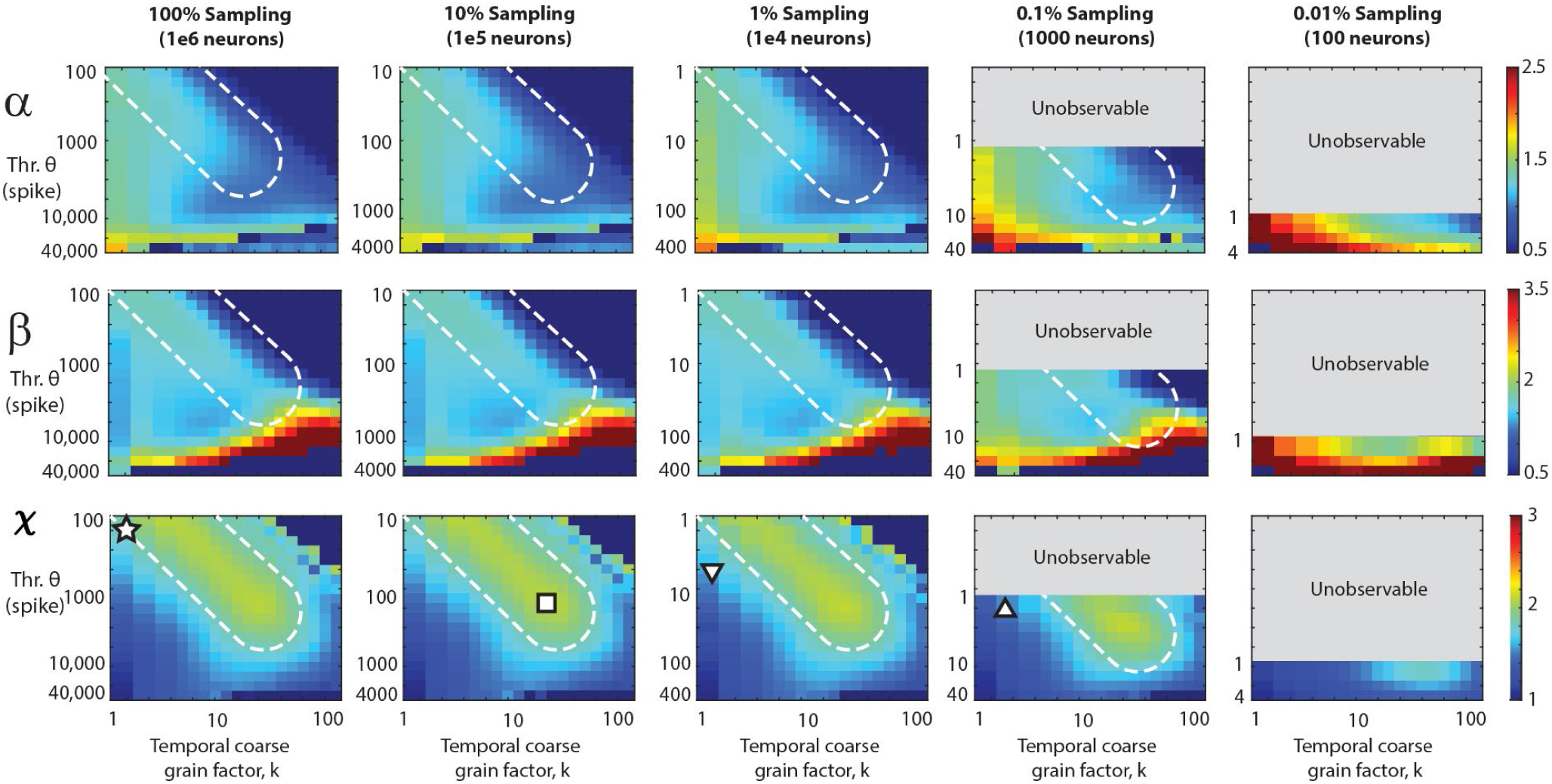
Consolidated view of the exponents α, **β** and ***X*** as a function of threshold and temporal coarse-graining for different sampling fractions. Consolidated view of α, β and *χ* as a function of *θ* and *k* for different values of *f*. For *f* = 100%, *χ* ≅ 2 for low *θ* and *k* (*star*) but as we make the data sparser by increasing the threshold (or reducing the sampling), we need a higher coarse graining factor, *k*, to compensate and rescue *χ* back to 2 (*square*). Bottom row is replotted for *χ* from Figure 4 for ease of comparison with the corresponding size slopes, *α*, and duration slopes, *β*. *White dotted region*: visual guide for *χ* close to 2 for increasing *θ* and *k*. The grey parts of the plots for f = 0.1% and 0.01% respectively are unobservable parameter regions since they would require fractional thresholds, below the 1 spike minimum resolution of the model (*cf.* Figure 4).

